# Antibiofilm agents with therapeutic potential against enteroaggregative *Escherichia coli*

**DOI:** 10.1101/2021.11.05.467448

**Authors:** David A. Kwasi, Chinedum P. Babalola, Olujide O. Olubiyi, Jennifer Hoffmann, Ikemefuna C. Uzochukwu, Iruka N. Okeke

## Abstract

**Background:** Enteroaggregative *Escherichia coli* (EAEC) is a predominant but neglected enteric pathogen implicated in infantile diarrhoea and nutrient malabsorption. There are no non-antibiotic approaches to dealing with persistent infection by these exceptional colonizers, which form copious biofilms. We screened the Medicines for Malaria Venture Pathogen Box for chemical entities that inhibit EAEC biofilm formation.

**Methodology:** We used two EAEC strains, 042 and MND005E, in a medium-throughput crystal violet-based antibiofilm screen. Hits were confirmed in concentration-dependence, growth kinetic and time course assays and activity spectra were determined against a panel of genome-sequenced EAEC. Antibiofilm activity against isogenic EAEC mutants, molecular docking simulations and comparative genomic analysis were used to identify the mechanism of action of one hit.

**Principal findings:** In all, five compounds (1.25%) reproducibly inhibited biofilm accumulation by at least one strain by 30-85% while inhibiting growth by under 10%. Hits exhibited at least 10-fold greater antibiofilm activity than nitazoxanide, the only known EAEC biofilm inhibitor. Reflective of known EAEC heterogeneity, only one hit was active against both screen isolates, but three hits showed broad antibiofilm activity against a larger panel of strains. Mechanism of action studies point to the EAEC anti-aggregation protein (Aap), dispersin, as the target of compound MMV687800.

**Conclusions:** This study identified five compounds not previously described as anti-adhesins or Gram-negative antibacterials with significant and specific EAEC antibiofilm activity. One molecule, MMV687800, targets the EAEC Aap. *In vitro* small-molecule inhibition of EAEC colonization opens a way to new therapeutic approaches to preventing and treating EAEC infection.

**Author summary:** Diarrhoea accounts for over half a million deaths in children under five annually. It additionally contributes to childhood malnutrition as well as growth and development deficiencies, particularly in low-income countries. Enteroaggregative *Escherichia coli* (EAEC) causes diarrhoea that is often persistent and can also contribute to growth deficiencies in young children. EAEC is a neglected pathogen that is often resistant to antimicrobial drugs. Small molecules that block EAEC colonization may hold the key to interfering with EAEC disease without promoting antimicrobial resistance. We screened the Medicines for Malaria Ventures Pathogen Box for chemicals that can interfere with EAEC biofilm formation, a key colonization indicator. Our screen identified five biofilm-inhibiting molecules that did not interfere with bacterial viability and therefore are unlikely to exert strong pressure for resistance. Molecular biology and computational investigations point to the EAEC anti-aggregative protein, also known as dispersin, as a possible target for one of these hit molecules. Optimizing EAEC antibiofilm hits will create templates that can be employed for resolving EAEC diarrhoea and related infections.

## Introduction

Diarrhoea constitutes a huge global disease burden [1,2], accounting for approximately 500,000 deaths among under-fives annually [3, 4]. Diarrhoea also contributes significantly to malnutrition as well as growth and development shortfalls, particularly in low-income countries [5,6]. The burden from diarrhoea is highest in Africa with Nigeria topping the list on the Africa continent and ranking second only to India’s contribution to the global burden from the syndrome [4].

Infectious diarrhoea can be caused by a wide range of micro-organisms including, but in no way limited to rotavirus, astrovirus, norovirus, *Entamoeba, Cryptosporidium*, multiple subtypes of diarrhoeagenic *Escherichia coli* and *Salmonella* [7]. Enteroaggregative *Escherichia coli* (EAEC) is a diarrhoeagenic *E. coli* subtype known to cause both acute and persistent diarrhoea (the latter continuing for more than 14 days) [8]. EAEC strains are additionally implicated in traveler’s diarrhoea and foodborne outbreaks worldwide [9-11]. EAEC are epidemiologically important globally and are repeatedly detected at high prevalence in many epidemiological studies [12-14], including our earlier and ongoing research in West Africa [15-18]. The burden of EAEC infections and their impact on child health necessitates a clear understanding of the pathogenesis of the disease, as well as effective interventions. However, EAEC research is neglected even more than research on other high-burden bacterial diarrhoeal pathogens such as rotavirus, enterotoxigenic *E. coli* and *Shigella* [19,20].

Hallmarks of EAEC infection include copious adherence to epithelial cells in a striking ‘stacked brick’ or ‘aggregative’ fashion as well as the formation of voluminous biofilms [16,21,22]. Biofilms are a complex community of organisms encased in an extracellular matrix and EAEC *in vitro* and *in vivo* biofilms are believed to contribute to persistent colonization and transmission of diarrhoea [23-25].

EAEC are genetically heterogeneous and difficult to delineate from commensals [20, 26, 27]. Although the molecular epidemiology of EAEC infection remains unclear, most strains colonize the intestinal mucosa via the aggregative adherence fimbriae (AAFs) and other non-structural adhesins [25,28], which also contribute to copious biofilm formation and subsequent host pathogenesis [23,27,29]. These critical components of the EAEC pathogenic cascade suggest that molecules which can interfere with or inhibit the assembly or function of adhesins central to EAEC virulence could be promising therapeutic candidates.

We hypothesized that small molecules can interfere with or inhibit EAEC adherence to host cells either by structurally modifying adhesins or competing for their receptor sites without inhibiting growth. We tested our hypothesis by screening the Medicines for Malaria Ventures’ (MMV) Pathogen Box chemical library (https://www.mmv.org/mmv-open/pathogen-box/about-pathogen-box), deploying a commonly used biofilm assay protocol [28,30-32], adapted to medium-throughput. Pathogen Box is a curated compound library containing 400 synthetic compounds arrayed in a 96-well format. Information about activities of the compounds against *Plasmodium falciparium* and some neglected human pathogens is provided with the library, but the compounds have not been tested against bacterial causes of diarrhoea or other Gram-negative bacteria. The structures, physicochemical properties as well as preliminary profiles of the compounds are provided with the box, providing useful insights and good chemical starting points for neglected pathogen drug research.

## Methods

### Bacteria strains, plasmids, and culture conditions

Two EAEC strains, 042, a prototypical EAEC strain originally isolated in Peru [33,34] and MND005E, an EAEC strain isolated from an ongoing case-control study in our laboratory in Nigeria [18], were used for the preliminary screens and confirmatory assays. These strains and 25 other biofilm-forming EAEC strains (Table 1) are isolates from diarrhoea patients under the age of five. Isogenic 042 mutants and laboratory strain ER2523 (NEB express) shown in Table 1 were used to obtain preliminary information on mode of action of hits. Molecular microbiology methods, comparative genomics and molecular docking were used to confirm hit mechanism of action. Strains were routinely cultured in Luria Bertani (LB) broth (Sigma Aldrich: Cat no. L3522), adding ampicillin (100 µg/ml) where necessary. Strains were archived at -80°C in sterile cryogenic vials in Luria Bertani broth (Mueller) glycerol in ratio 1:1. Plasmids used or generated in the study are listed in Table 1.

**Table 1:**
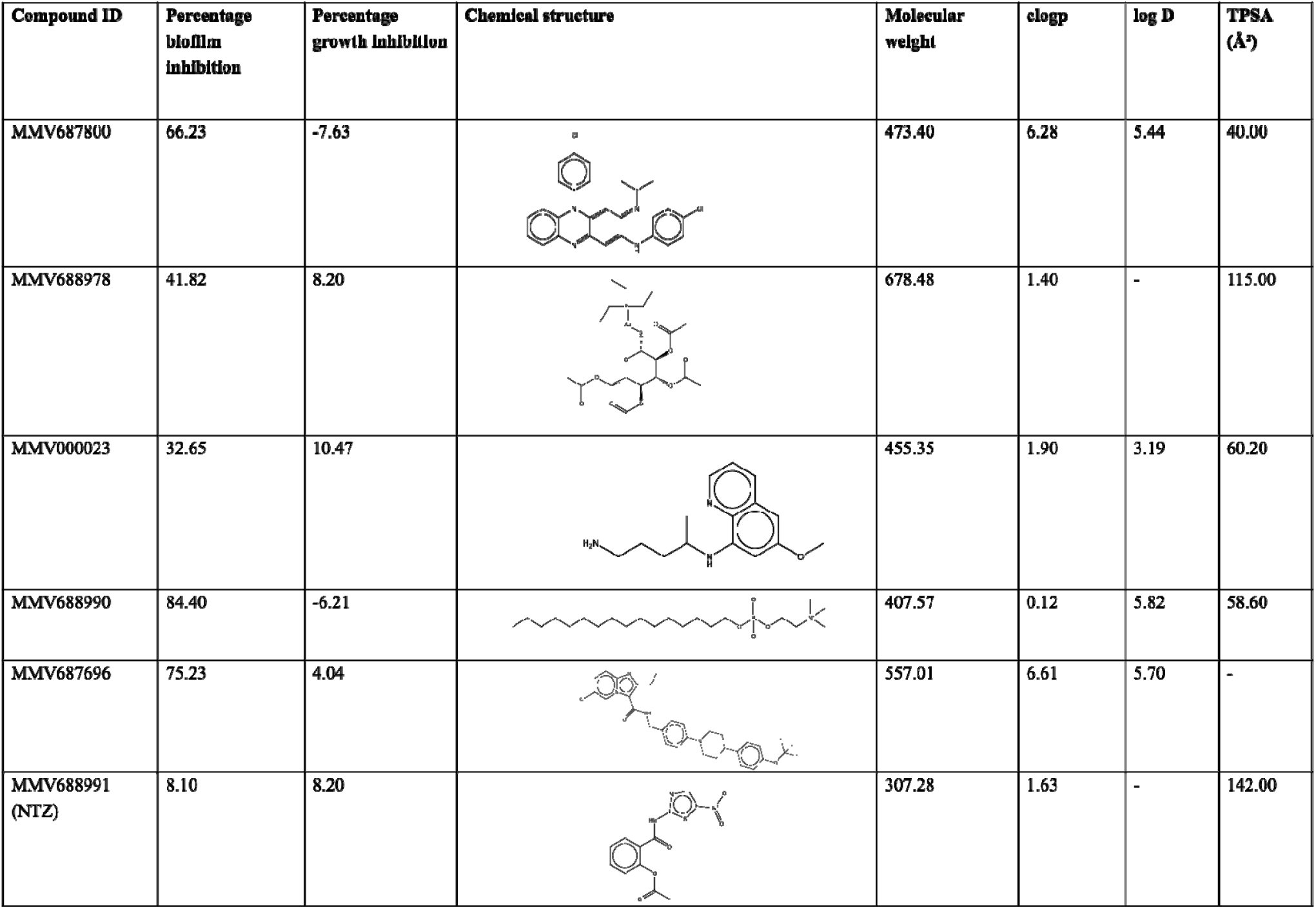
Chemistry of validated hits: structures, and molecular descriptors. Data provided by MMV and PubChem.

| Compound ID | Percentage biofilm inhibition | Percentage growth inhibition | Chemical structure | Molecular weight | clogp | log D | TPSA (Å²) |
| --- | --- | --- | --- | --- | --- | --- | --- |
| MMV687800 | 66.23 | -7.63 |  | 473.40 | 6.28 | 5.44 | 40.00 |
| MMV688978 | 41.82 | 8.20 |  | 678.48 | 1.40 | - | 115.00 |
| MMV000023 | 32.65 | 10.47 |  | 455.35 | 1.90 | 3.19 | 60.20 |
| MMV688990 | 84.40 | -6.21 |  | 407.57 | 0.12 | 5.82 | 58.60 |
| MMV687696 | 75.23 | 4.04 |  | 557.01 | 6.61 | 5.70 | - |
| MMV688991 (NTZ) | 8.10 | 8.20 |  | 307.28 | 1.63 | - | 142.00 |

### Chemical library

Pathogen Box, a chemical library of 400 diverse, drug-like molecules, was a kind gift from the Medicines for Malaria Venture (MMV, Switzerland). Compounds were supplied at 10 mM dissolved in 10 µl dimethyl sulfoxide (DMSO) in sterile 96-well polystyrene plates (A-E, 80 compounds per plate). Tentative hits were resupplied compounds from MMV and/or purchased from Sigma Aldrich for downstream experiments. These were received as powders then reconstituted in our laboratory in DMSO (Sigma Aldrich: Cat no. D5879). In every set up, final concentration of DMSO did not exceed 1%, which we confirmed in preliminary assays had no effect on EAEC growth and biofilms. Stock solutions were stored at -20°C, thawed, mixed and then diluted to required concentrations prior to each experimental set up.

### Biofilm inhibition assays

The medium throughput screen was set up in sterile 96-well flat bottom polystyrene plates (Nunc: 260860). For each assay, 5 µl of 200 µM drug solutions dissolved in net DMSO (Sigma Aldrich: Cat no. D5879) were pipetted into the 96-well plate using a multichannel pipette (Gilson). Assay plates subsequently received 195 µl of high glucose Dulbecco’s Modified Eagles Medium (DMEM, ThermoFisher Scientific, cat no. 11965092) containing overnight culture in ratio 1:100 to achieve a final drug concentration of 5 µM in each test well. Control wells received 2 µl of DMSO vehicle and 198 µl of the same mixture of DMEM and overnight culture to ensure a final DMSO concentration of 1% in control wells [35]. Assay was set up in triplicate for control and each compound then incubated at 37°C for 8 h. Planktonic cell growth was determined by quantifying optical densities at 595 nm using a microplate spectrophotometer (Thermo scientific multiscan FC: Cat no. 51119000). Plates were thoroughly washed three times with 200 µl of sterile PBS per well using a microplate washer (Micro wash 1100 from Global diagnostics) then air-dried and fixed with 75% ethanol for 10 minutes. After the plates were dried, we stained with 0.5% crystal violet for 5 minutes then washed thoroughly with water. We dried plates completely then eluted crystal violet using 200 µl of 95% ethanol for 20 minutes. Biofilm was quantified by determining optical density of eluted crystal violet at 570 nm using a multiscan microplate spectrophotometer.

Antibiofilm and growth inhibitory effects of each compound were computed from the averages of 3 replicates applying the following formulae:

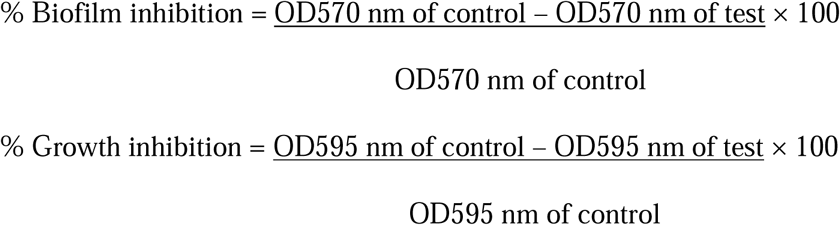

Antibiofilm activity of compounds identified as initial hits was additionally confirmed in concentration dependent assays set up in sterile flat bottom 96-well polystyrene plate. Concentrations of compounds were prepared in serial 2-fold dilutions ranging from 0.3125 µM to 20 µM. Serially diluted compounds were added in triplicate to 195 µL of DMEM containing standardized overnight culture of organism. For retesting of previously described antibiofilm agent nitoxazanide (NTZ); 15µg/mL (48.8µM), 20 µg/mL (65.15µM) and 25 µg/mL (81.45 µM) earlier reported to exhibit significant biofilm inhibition in EAEC [28] were tested.

Plates were incubated at 37°C for 8 h and growth determined at 595 nm. Plates were washed, fixed, and stained after which biofilm (eluded crystal violet) was determined at 570 nm. Time-course assays were set up independently in triplicate using drug concentrations prepared in serial 2-fold dilutions ranging from 0.3125 µM to 20 µM at time points 4, 8, 12, 18, 24 and 48 h respectively. To determine antibiofilm spectra, 2 reference strains (042 and 60A), 25 EAEC strains identified from cases of childhood diarrhoea in an on-going study in Nigeria [18,33] were used and isogenic EAEC 042 mutants (Table 1) were used to infer mechanisms of action involving known surface factors.

### Growth kinetic assay

We monitored growth in the presence of varied concentrations of hit compounds at 37°C in sterile 96-well polystyrene plates. Concentrations of compounds were prepared in serial 2-fold dilutions ranging from 0.3125 µM to 20 µM in DMEM. Thereafter, serially diluted compounds were added, assay was set in triplicate for each hit concentration and optical density was determined at 595 nm using a microplate spectrophotometer at 0 mins, 30mins, then hourly up until the 8^th^ hour, incubating at 37°C between readings.

### Molecular docking

In order to obtain atomic level insight into and to validate the mechanism of action of the identified hits, we employed a molecular docking protocol that screened the identified hit molecules against the anti-aggregation protein (Aap) [36], the likely antibiofilm target from our mutant and compliment testing assays. The employed Aap structure (accession code 2JVU.pdb [36]), solved using solution NMR, comprises 20 structurally distinct conformers all of which were individually employed as receptor molecules, a strategy that allowed for the incorporation of macromolecular flexibility in the docking protocol. For the docking screening, three dimensional models were first generated for the five identified MMV molecules using the ChemBioOffice Suite. Energy minimization with the steepest descent algorithm was then performed on each model to resolve steric clashes and identify each molecule’s potential energy minimum from the sampled conformational landscape. Each model was then saved in the P rotein D ata B ank (PDB) format.

Gasteiger atomic charges, needed for a more accurate computation of electrostatics of the binary interactions, were calculated using AutoDock Tool [37,38] for each of the five MMV hits as well as for each of the twenty Aap structures. AutoDock was also employed in setting up a hyperrectangular docking grid with x, y, z dimension (in Ang strom units) of 47.25, 36.0, 36.0 centred at 72.234, 0.223, 1.139, respectively. The grid dimensions were carefully selected to achieve a total coverage of the Aap models. Using AutoDock Vina [39], each of the five hit compounds was subsequently subjected to docking screening against each of the twenty Aap conformers using a rigid docking protocol with bonded degrees of freedom DOFs) frozen in the receptor macromolecules while all torsional DOFs in the five hit compounds were geometrically optimized during conformational search for stable ligand-receptor complexes. The best binding free energies computed for each of the five hits from this ensemble docking approach were then averaged over the twenty Aap conformations.

A similar docking was performed for subunits of the aggregative adherence fimbriae variant II (AAF/II), a previously reported antibiofilm targeted in EAEC 042 by nitazoxanide [28]; the AAF/II is suspected as a second target for one of the hits from the outcome of the 042 mutant testing. The solution NMR spectroscopy structure of AAF/II major subunit (accession code 2MPV [8]), and the dimerized crystallographic (3.0 Å) AAF/II minor subunit (accession code 4OR1 [8]) were retrieved from www.rcsb.org. The five hits and NTZ were docked against the macromolecular structure of the AAF/II using docking grid with xyz dimensions (unit of Å) of 12.890, 11.530, 11.697 and centred at 13.002, 1.147, -9.750 for the AAF/II major subunit. In the case of the minor subunits xyz dimensions of 8.765, 11.831, 11.142 and centred at -22.572, 51.865, -2.643 were employed.

### Complementing the *aap* mutant

PCR primers were designed to amplify the *aap* (antiaggregative protein) gene from 042 genomic DNA. The sequence of primers used were: GGC<u>caattg</u>atgaaaaaaattaagtttgttatcttttc and GATC<u>CCTGCAGG</u>ttatttaacccattcggttagag with *Mfe*I (Cat. No. R0589S) and *Sbf*I (Cat. No. R0642S) restriction enzyme recognition sites incorporated into the tails respectively for cloning into pMAL-c5X vector (NEB, Cat. No. N8018S). The *aap* gene was amplified using the following cycle: 94°C for 2 min, then 25 cycles of 94°C for 30 sec, 55 °C for 45 sec and 72°C for 30 sec followed by a final extension of 72°C for 15 mins. PCR products were purified using Zymo DNA Clean & Concentrator™-5 Kit (Cat. No. D4003), digested with *Mfe*I and *Sbf*I, and ligated into pMAL-c5X vector using T4 ligase (Cat. No. M0202S). The recombinant plasmid obtained, was cloned into ER2523 (NEB express, Cat. No. E4131S) and resistant clones were isolated on LB-ampicillin plates (100 µg/ml). The resulting clone, pDAK24, was verified using PCR to confirm the presence of the *aap* insert in plasmid, extracted using the QIAprep^®^ Spin Miniprep Kit (Cat. No. 27106), and sequenced. The verified pDAK24 clone was used to transform LV1 (042Δ*aap*), LV2 (042Δ*aap*Δ*hra1*) *and* LTW1 (042Δ*aap*Δ*aafA*) [40] and the resulting *aap* mutant complements were tested for biofilm formation and inhibition.

### Comparative analysis of virulence genes

Antibiofilm spectra of the five hits identified in this study were investigated using EAEC reference strain 042 [33], 60A from Mexico [41] and 25 other EAEC strains identified and characterized in our laboratory in Nigeria from an ongoing diarrhoea case-control study [18]. Virulence gene profiles of strains in each group were retrieved from whole genome sequence data using VirulenceFinder database [42] then compared to identify genes which were unique to each group for each hit.

### Statistical analysis

Data from biofilm inhibition assays (average of three replicates) were analyzed by comparing inhibition / percentage inhibitions for tests and controls and significant differences were inferred from Fisher’s exact test (to identify genes which are unique to strains whose biofilms were inhibited by hits) and student’s t-test analysis (for biofilm inhibition assay).

## Results

### Pathogen Box contains compounds capable of inhibiting EAEC biofilm formation

The 400-compound Pathogen Box chemical library was screened for EAEC biofilm formation inhibitors in a medium throughput assay. Initial hit selection criteria were that molecules must demonstrate at least 30% biofilm inhibition and under 10% growth inhibition at a concentration of 5 µM [43,44]. Applying these criteria, we identified five compounds which reproducibly inhibited accumulation of biofilm biomass by EAEC 042 and/ or MND005E by 30-85% while inhibiting growth by ≤ 10% (Figure 1a-d). As shown in Table 1, the compounds possessed diverse molecular architectures and chemical properties. Consistent with the known heterogeneity of EAEC strains, the two EAEC test strains, which were selected because they were phylogenetically distant and had largely non-overlapping suites of virulence factors, retrieved different hits from the library. Compounds MMV687800, MMV688978 and MMV687696 inhibited biofilm formation of strain 042 only, MMV000023, was active against MND005E while MMV688990 showed activity against both strains. Nitazoxanide (MMV688991), the only previously reported EACE 042 biofilm inhibitor [28], is also contained within Pathogen Box but did not meet our hit criteria.

**Figure 1:**
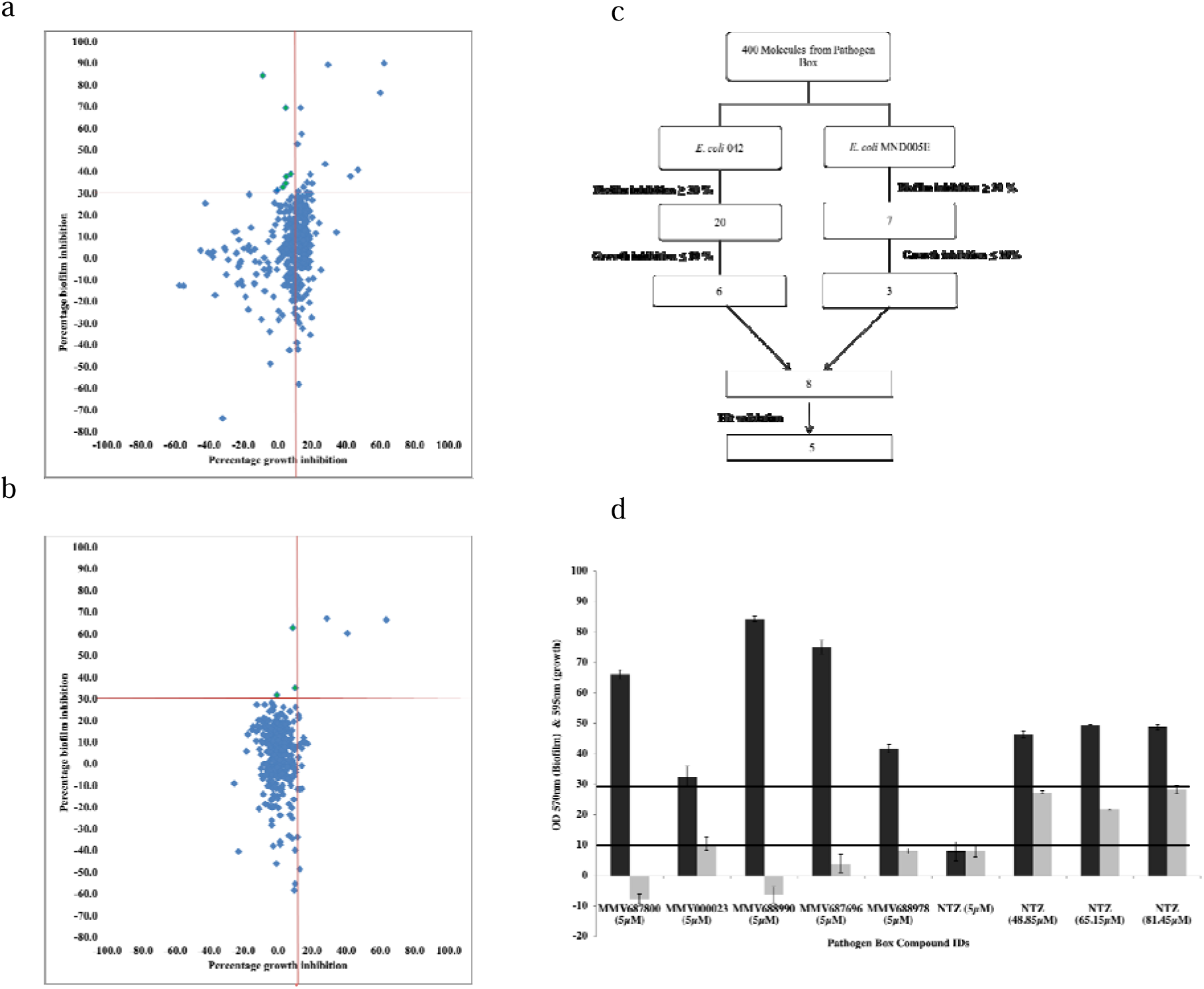
(a-b) Preliminary screen outcome of 400 drug-like molecules in Pathogen Box as a measure of percentage biofilm inhibition against growth inhibitions at 5µM for (a) EAEC strain 042 and (b) EAEC strain MND500E. Each compound except the antibiofilm hits is represented by a blue diamond. Compounds in the top portion of the scatterplot (above the horizontal red line) show >30% biofilm inhibition activity and those to the left of the vertical line additionally inhibit growth by 10%. The number of initial hit candidates (colored green) was six for 042 in (a) and three for MND005E in (b). (c) Hit progression cascade leading to 5 validated EAEC biofilm non growth inhibiting compounds. A total of 8 hits were obtained (One hit inhibited biofilms without inhibiting growth in both strains), but only five (validated hits) reproducibly inhibit biofilms in EAEC. (d) Biofilm inhibition (black bars) and growth inhibition (grey bars) of hit compounds and nitazoxanide against (042 and MND005E). The five hits inhibited biofilm formation by over 30% (upper horizontal line) while inhibiting growth by under 10% (lower line). NTZ showed activity but not within the range of these

Significant antibiofilm activity was previously reported for NTZ at 15 µg/ml (48.8µM), 20 µg/ml (65.15µM) and 25 µg/ml (81.45µM) with an estimated growth inhibition of up 50% [28]. Based on this information, we conducted comparative biofilm inhibition assays for our hits (at 5 µM) and NTZ at 48.8 µM, 65.15 µM and 81.45 µM. We observed similar patterns of antibiofilm activity and growth inhibition with NTZ at these concentrations as reported earlier by Shamir et al. [28]. Additionally, the five validated hits in this screen were found active at concentrations at least 10 times lower than those of nitazoxanide (NTZ) as shown in Figure 1c. Consequently, hits from this screen are potent biofilm inhibitors and we cannot rule out the presence of other weak biofilm inhibitors in Pathogen Box (Figure 1a-b). The class (medicinal indication) of the 400 Pathogen Box compounds matched with biofilm inhibition outcome is summarized in Table S1. Four of the five validated hits are reference compounds with other known activities, while one is indicated for tuberculosis. The three unvalidated hits (which were not retrieved consistently in confirmatory assays especially in concentration dependent and growth kinetics assays applying our hit selection criteria) in this study are kinetoplastid inhibitors.

In the course of the screen, we observed that known antibacterials in Pathogen Box, doxycycline, levofloxacin and rifampicin inhibited biofilm formation by 60.33, 89.92, and 90.08% respectively but also inhibited growth by 15.07, 64.53 and 32.53. Our initial preliminary screens additionally turned out MMV688362, MMV688771 and MMV676159 as EAEC 042 hits. However repeated testing and further validation in confirmatory assays revealed antibiofilm activity were due to antibacterial activity in one case, with the others exhibiting a little lower antibiofilm activity than the hit criteria minimum cut off (30 %). Five additional compounds also inhibited both growth and biofilm formation and are being further evaluated as potential antibacterial hits.

Planktonic cell growth of EAEC 042 and MND005E in the presence of 0.3125 – 20 µM of hits over a time course of 0 to 8 h was not significantly different from the compound-free control (Figure 2).

**Figure 2.**
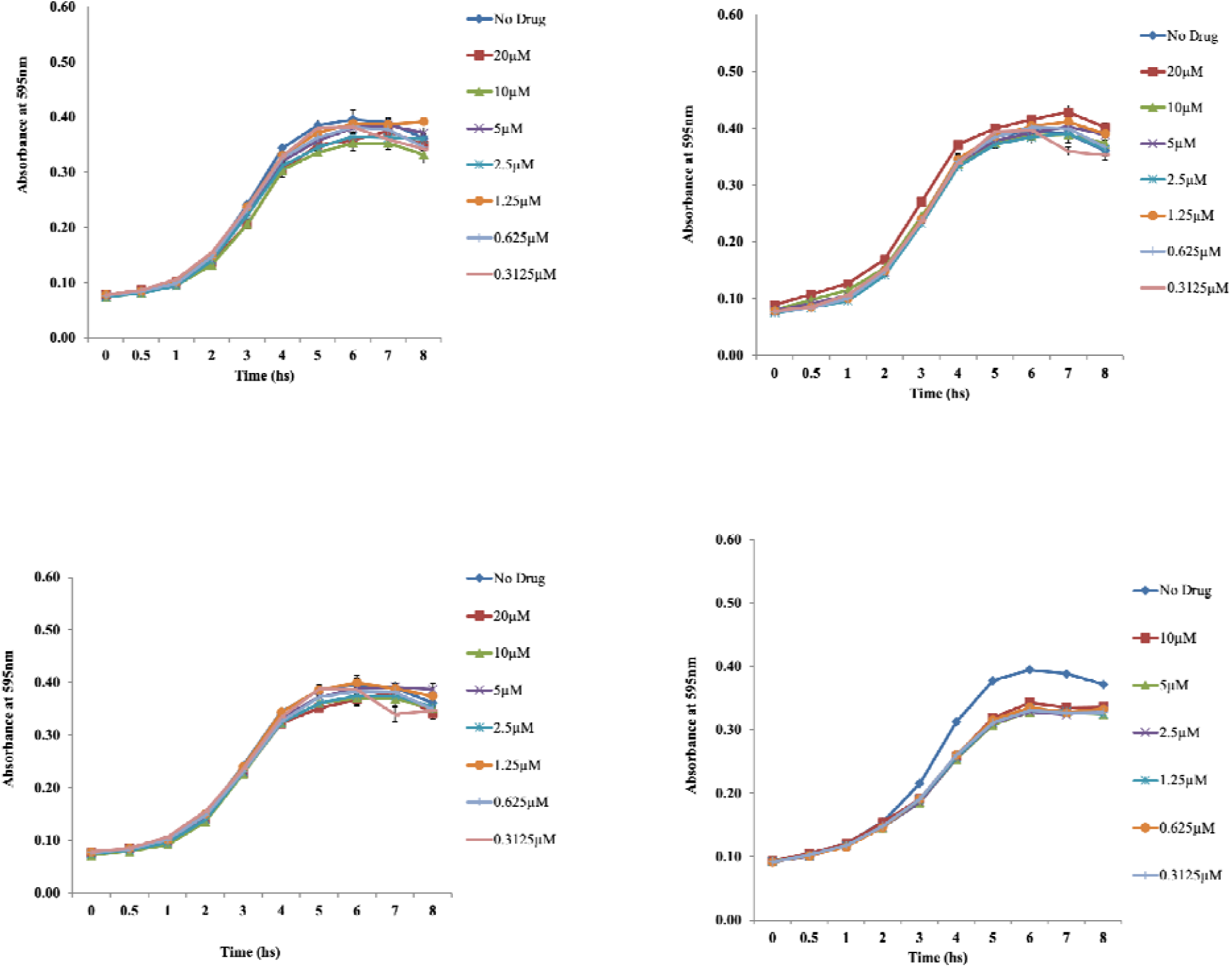
Hits did not significantly alter growth kinetics of EAEC strains at the various concentrations tested over 8 h: Growth kinetics of EAEC 042 with different concentrations of **a**) MMV688978, b) MMV687800, c) MMV688990, d) Growth kinetics of MND005E with different concentrations of MMV688990.

We tested concentration-dependency of three of our five hits, MMV687800, MMV688978 and MMV688990 against EAEC strain 042. As shown in Figure 3, the three compounds exhibited concentration-dependent inhibitory effects on biofilm formation in EAEC 042 with correlations (r^2^) of 0.9552, 0.9355 and 0.9593 for hit MMV688978, MMV687800, MMV688990 respectively. Individual analysis of concentration-dependent curves revealed that hits exhibited potent activity, with concentrations as low as 2.5 µM inhibiting biofilm formation by up to 30 % (Figure 3).

**Figure 3:**
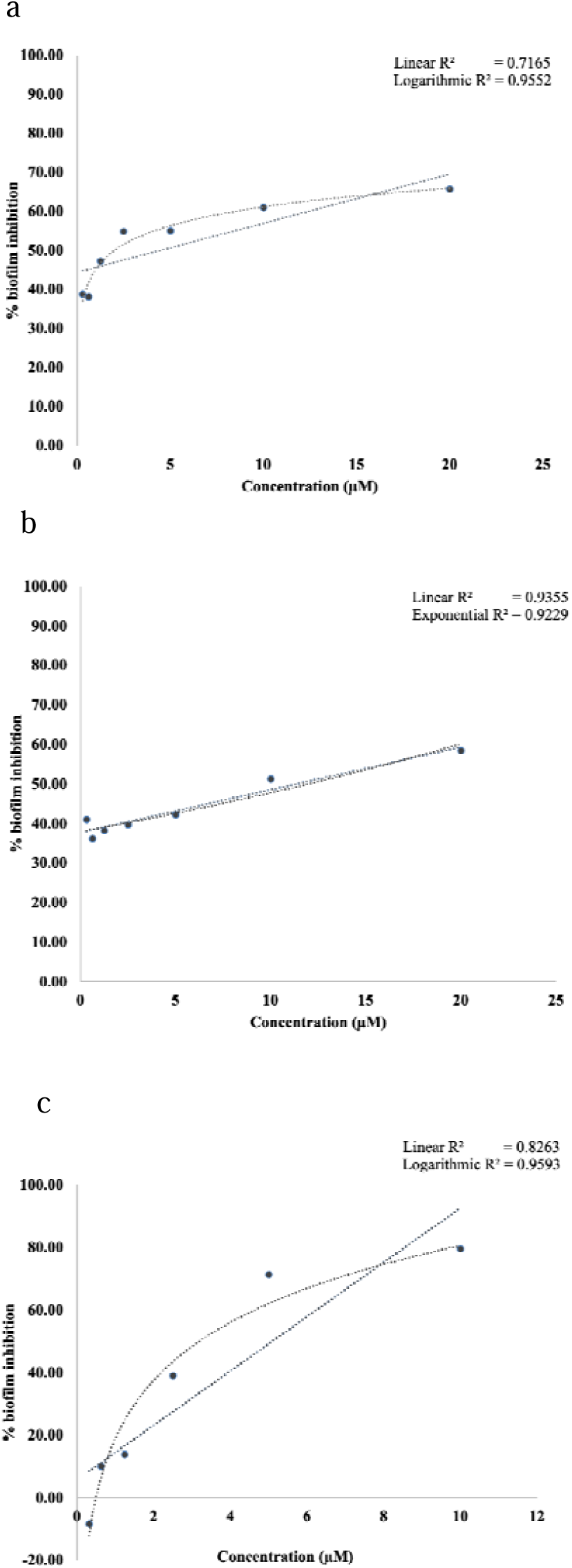
Regression analysis showing the relationship between percentage biofilm inhibition and drug concentrations by: a) MMV688978, (b) MMV687800, (c) MMV688990. Hits demonstrated concentration dependent biofilm inhibition and were active (meeting >30 % biofilm inhibition cut-off) at concentrations as low as 2.5 µM.

When we tested inhibition of biofilm formation across a 48 h time-course, we found that the biofilm inhibition effect was most pronounced at early time points and then again after the biofilm was well established (Figure 4) and (Figure S1).

**Figure 4:**
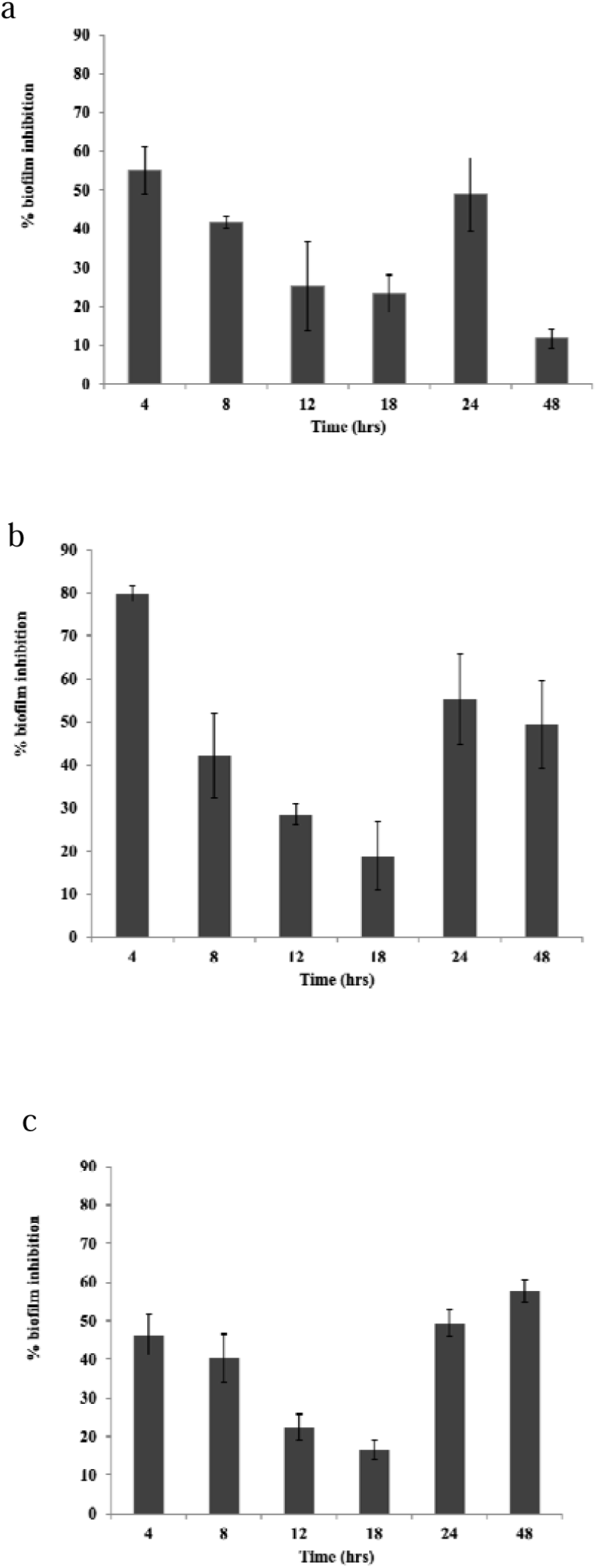
Time course assay of 042 biofilm (black) by 3 hits at 5 µM shows inhibition sits ahtighe the earlier time points then at later phase in: (a) MMV688978 (b) MMV687800 (c) MMV688990.

### Inhibition of biofilm formation in 042 *aap, hra1* and *aaf* mutants

As a first step towards identifying mechanisms of action, we tested biofilm inhibition in single and double mutants in three key colonization genes of EAEC strain 042 against the four hits that inhibit biofilm formation in this strain. MMV687696, MMV688978 and MMV688990 significantly inhibited biofilm formation by all five mutants at a level similar to wildtype 042 inhibition (Figure 5b-d). Noteworthily, MMV687800 reduced biofilm formation by 042Δ*hra1* (SB1) to a similar degree as the wildtype (EAEC 042) (Figure 5a), ruling out Hra1 as a target for this compound. A slightly lower degree of inhibition was seen in the 042Δ*aafA* mutant (3.4.14) but this was not statistically significantly different from the inhibition produced on the wild type (Figure 5a). For the *aap* mutant, 042Δ*aap* (LV1), inhibition occurred to a significantly lower degree. When we tested *aap* double mutants, we found that inhibition was similarly impaired in the 042Δ*aap*Δ*hra1* (LV2) and 042Δ*aap*Δ*aaf* (LTW1), with the latter showing no significant difference (p > 0.05) in biofilm formation in the presence or absence of MMV687800 (Figure 5a). Thus, the data point to *aap* as a likely target for MMV687800 and *aaf* may or may not be a minor target. Following the hypothesis that the antibiofilm activity of MMV687800 could involve *aap*, we set up biofilm assays with MMV687800 against EAEC 042, LV1, LV2, LTW1 alone and with the mutants complemented with pDAK24. As is known for these strains, biofilm inhibition was low in mutants 042Δ*aap*Δ*hra1* (LV2) and 042Δ*aap*Δ*aaf* (LTW1) double mutants compared to the wild type strain 042 (Figure 6). Conversely, notable reductions in biofilm formation were observed in the construct, LV1(pDAK24), LV2(pDAK24) and LTW1(pDAK24) (Figure 6), where *aap* deletions were complemented in *trans*.

**Figure 5:**
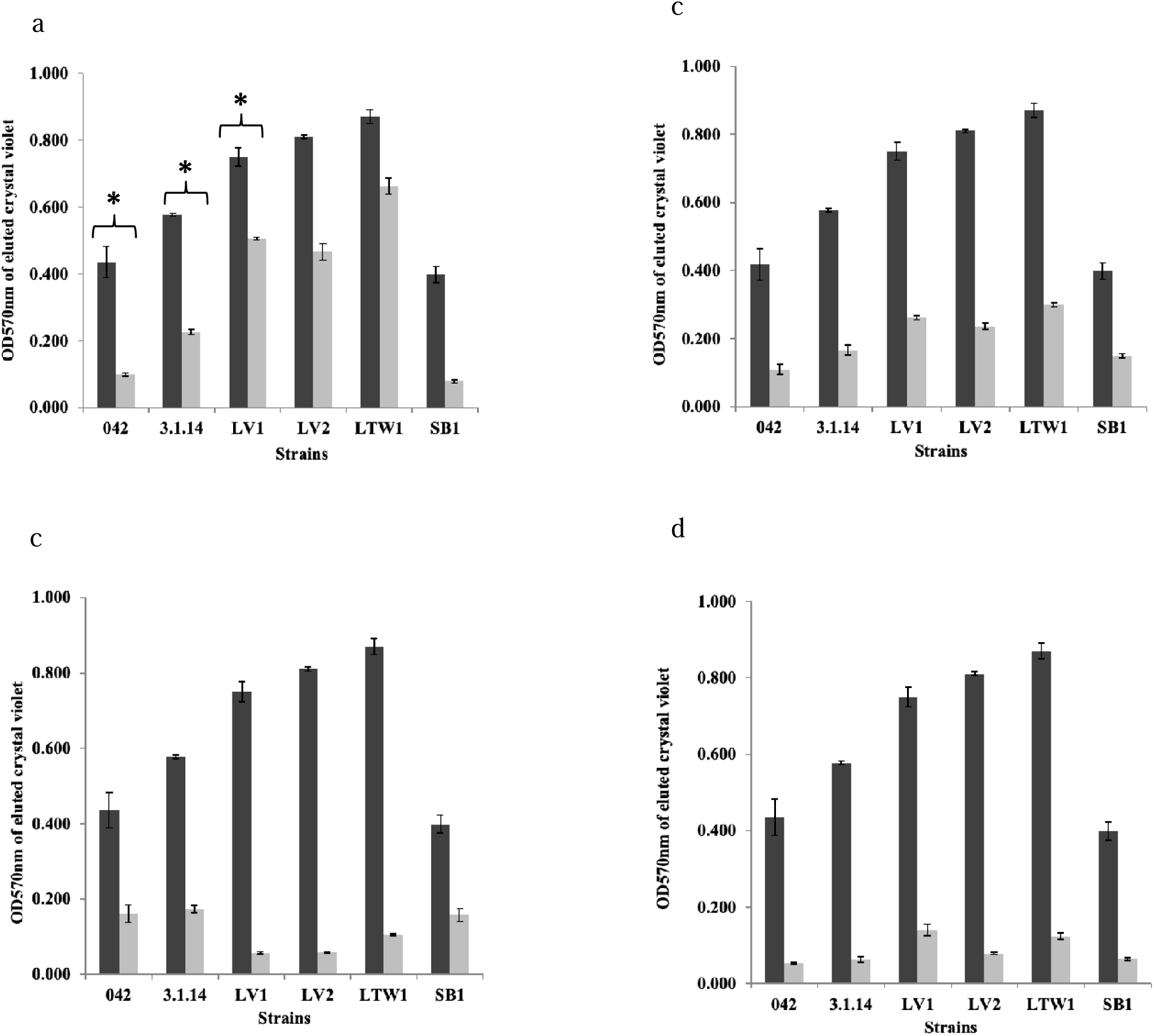
Effect of hits (grey) on biofilm formation (black) in *E. coli* 042 and 042 mutants: (a) MMV687800 (b) MMV687696 (c) MMV688978 (d) MMV688990. 042 is the prototypical EAEC strain from Peru, 3.1.14: 042Δ*aafA*, SB1: 042Δ*hra*, LV1: 042Δ*aap*, LV2; 042Δ*aap*Δ*hra1* and LTW1: 042Δ*aap*Δ*aafA*. Biofilm inhibition in wild type strain 042, and non-*aap* mutants 3.1.14 and SB1 was highly significant (^*^= P < 0.05), this was in contrast with inhibition in *aap* mutants LV1, LV2 and LWT1 (P > 0.05) indicating *aap* is a likely target for MMV687800.

**Figure 6:**
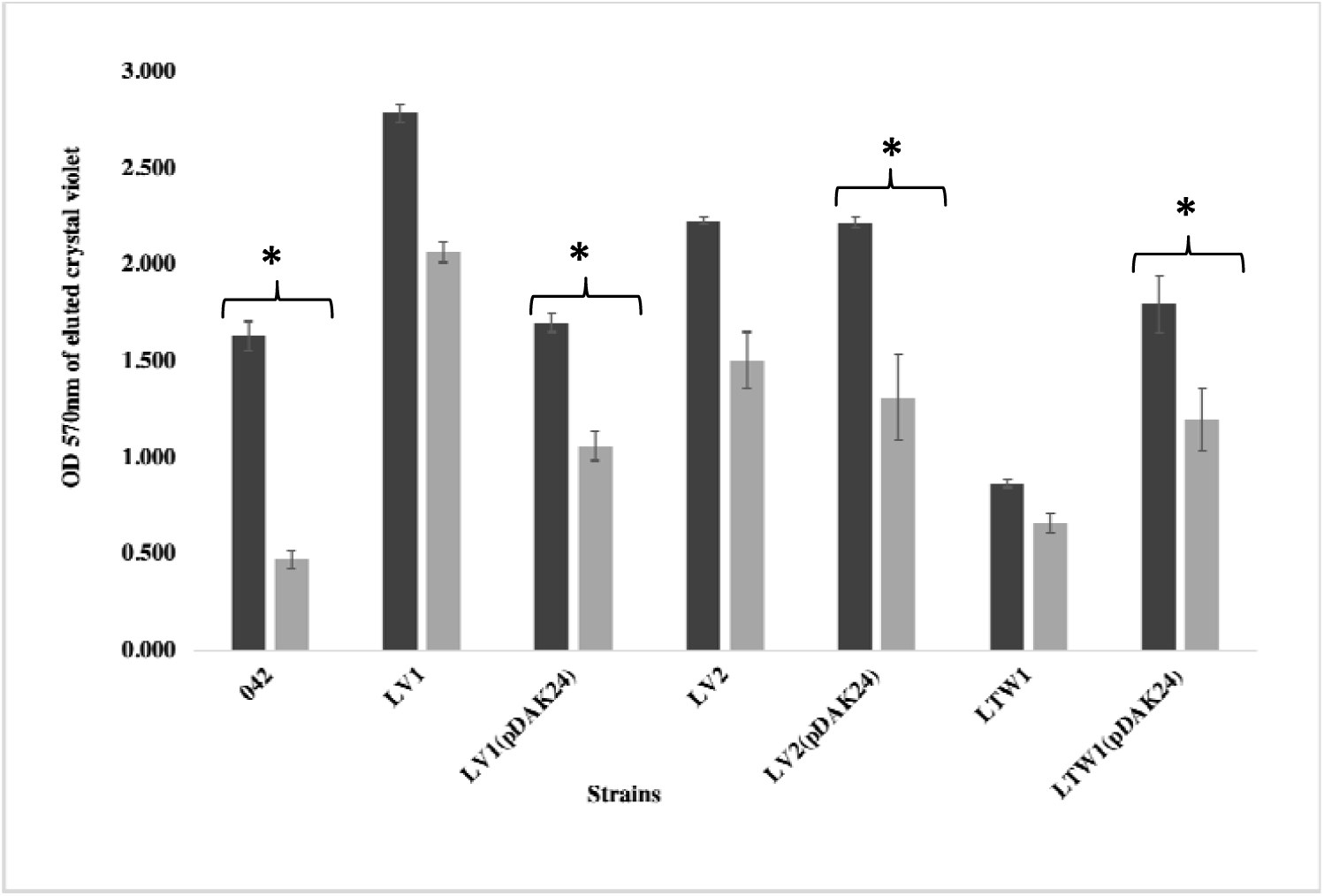
Biofilm inhibition of *E. coli* 042, 042 mutants and *aap* complimented strains by MMV687800. Biofilm formation (black) and inhibition by hit (grey). 042 is the prototypical EAEC strain from Peru, 3.1.14: 042Δ*aafA*, SB1: 042Δ*hra*, LV1: 042Δ*aap*, LV2; 042Δ*aap*Δ*hra1* and LTW1: 042Δ*aap*Δ*aafA*. LV1(pDAK24), LV2(pDAK24), and LTW1(pDAK24) are complement strains of LV1, LV2 and LTW1 mutants with *aap* inserts. Biofilm was significantly inhibited (^*^= P < 0.05) in wild type strain, and in *aap* complemented strains compared to *aap* mutants (P > 0.05). This fulfils Molecular Koch’s postulates and confirms *aap*’s involvement in biofilm inhibition by MMV687800.

Inhibition was not completely abrogated in *aap* mutants, hence it is probable that there is more than one MMV687800 target in 042 and collectively, the data suggest that *aafA* could represent a second target.

### Antibiofilm spectra of hit compounds

Along with the two EAEC strains used in our preliminary screen (EAEC strains 042 [33] and MND005E [18]), we investigated antibiofilm activity of hits against, 60A an EAEC isolate from Mexico [39.41] and 24 EAECs strains from our laboratory [18], which form moderate or strong biofilms and have been whole genome sequenced (Table 2). All five hits inhibited biofilms in different EAEC strains to varying degrees with MMV687800, MMV688978 and MMV688990 demonstrating broader antibiofilm activity spectra and likely different mechanisms of action (Table 3). Biofilm formation was inhibited by MMV687800 for all five isolates carrying allele 3 of the *aap* gene present in 042, but only four of 22 strains carrying other *aap* alleles or no *aap* gene (p=0.0016, Fisher’s exact test). Similarly, MMV687800 inhibited biofilms in all five strains bearing *aafA, aafB, aafC* and *aafD* genes but only four of 22 strains carrying none of the *aaf* genes (p = 0.0016, Fisher’s exact test).

**Table 2:**
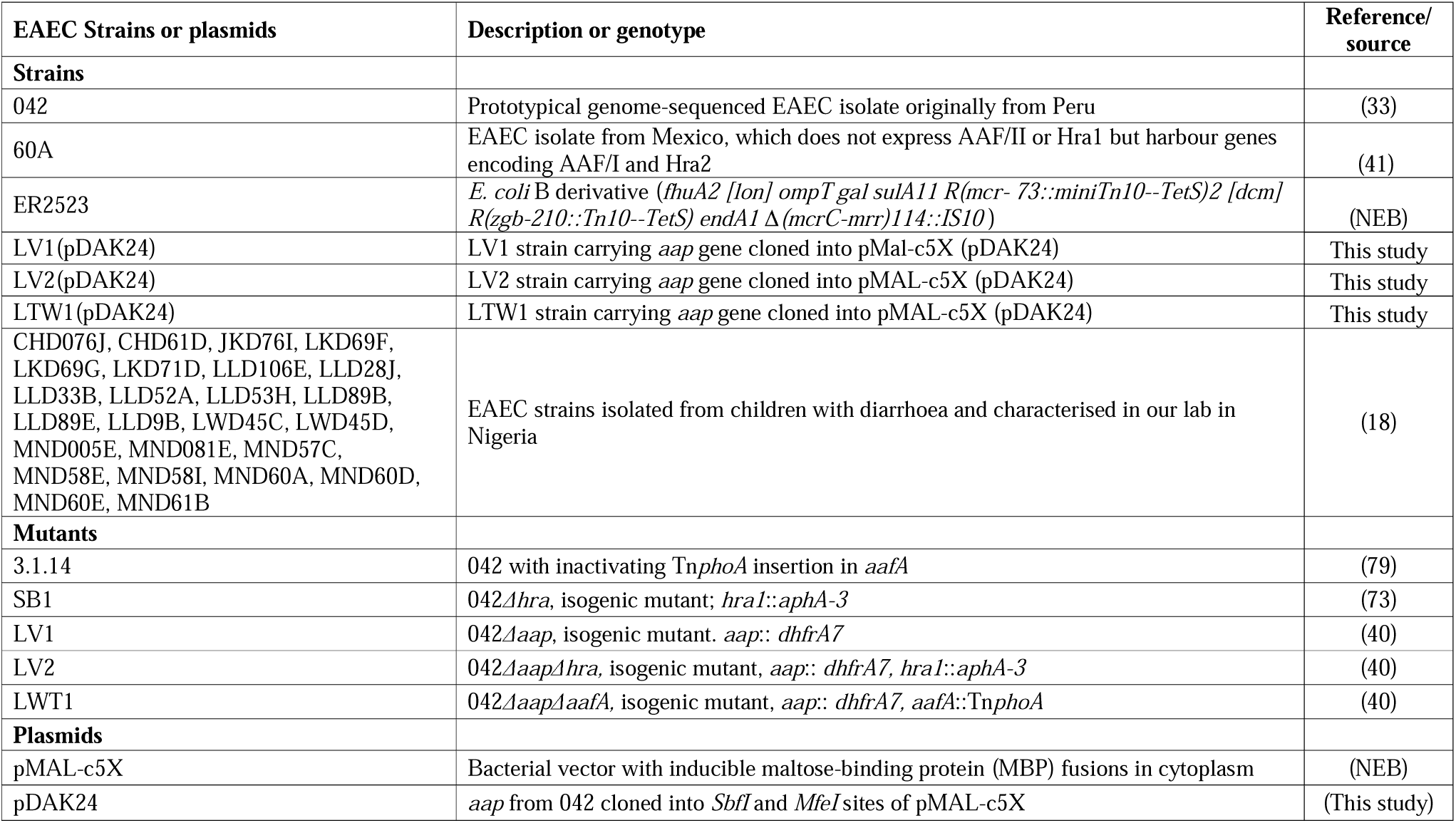
Reference strains and EAEC strains used in this screen.

**Table 3:** EAEC strains and antibiofilm activity spectra profile of 5 hit.

| Strains | Hits and percentage biofilm inhibition |  |  |  |  |
| --- | --- | --- | --- | --- | --- |
|  | MMV687800 | MMV688990 | MMV688978 | MMV678696 | MMV000023 |
| 042 | 76.52 | 69.87 | 60.70 | 47.19 | 7.20 |
| 60A | 35.60 | 59.31 | 39.02 | 38.73 | 61.25 |
| CHD076J | 5.85 | -18.46 | -12.62 | 13.69 | 0.31 |
| CHD61D | 18.94 | 12.52 | 24.28 | -3.88 | 3.79 |
| JKD76I | 45.82 | 12.50 | 25.15 | 4.85 | 20.22 |
| LKD69F | -21.59 | 1.06 | 27.12 | -15.50 | -48.34 |
| LKD69G | 6.23 | -9.86 | 12.46 | -16.26 | -69.27 |
| LKD71D | -6.15 | 13.20 | 3.39 | -53.39 | -3.47 |
| LLD106E | 77.28 | 38.06 | 47.08 | 45.06 | 35.01 |
| LLD28J | 52.02 | 51.27 | 59.80 | 15.54 | 58.09 |
| LLD33B | -10.88 | -5.88 | 1.47 | -19.12 | -16.76 |
| LLD52A | -50.74 | -19.37 | 18.53 | -32.56 | -46.46 |
| LLD53H | 44.15 | 19.84 | 48.12 | 43.43 | 37.09 |
| LLD89B | 11.64 | 57.28 | 39.84 | 33.27 | -20.39 |
| LLD89E | 1.69 | 6.26 | 36.15 | -14.61 | 32.00 |
| LLD9B | 7.68 | -15.45 | 4.25 | -11.47 | -3.89 |
| LWD45C | 60.86 | -17.73 | -13.03 | -11.99 | -12.38 |
| LWD45D | 45.19 | -28.02 | -16.80 | -15.18 | -15.75 |
| MND005E | -40.17 | 62.70 | 5.19 | -58.71 | 35.61 |
| MND081E | -1.39 | -14.49 | 17.96 | -35.00 | -33.99 |
| MND57C | 0.41 | -11.59 | 3.19 | -26.96 | -33.15 |
| MND58E | -11.59 | -21.77 | 2.96 | -1.35 | -11.59 |
| MND58I | -4.84 | -22.13 | -4.72 | -13.71 | -12.10 |
| MND60A | -50.80 | -9.83 | 9.48 | -32.47 | -40.95 |
| MND60D | -72.48 | -5.87 | 4.91 | -20.64 | -57.25 |
| MND60E | 36.00 | 15.89 | -2.02 | -13.00 | -47.09 |
| MND61B | -4.91 | -15.74 | 25.70 | -10.75 | -28.04 |

### Binding affinity of hit compounds with Aap and Aaf subunits

To validate Aap as a target for antibiofilm activity by MMV687800 and to determine whether AAF/II might be a target, we independently computed binding affinities of all five hits and NTZ (first reported to interfere with AAF/II and type I pili assembled by the chaperone usher pathway in EAEC) [28] with twenty Aap conformers and the AAF/II major and minor subunits using docking simulation. MMV687696 and MMV687800 demonstrated exceptionally strong interaction (in some cases ΔG ≤ -8.1 kcal/mol) with Aap as depicted in Figure 7. The other 3 hits exhibited comparatively lower affinity for Aap. As expected, (NTZ does not target Aap), there was also significantly lower affinity for Aap with NTZ (Figure 7).

**Figure 7:**
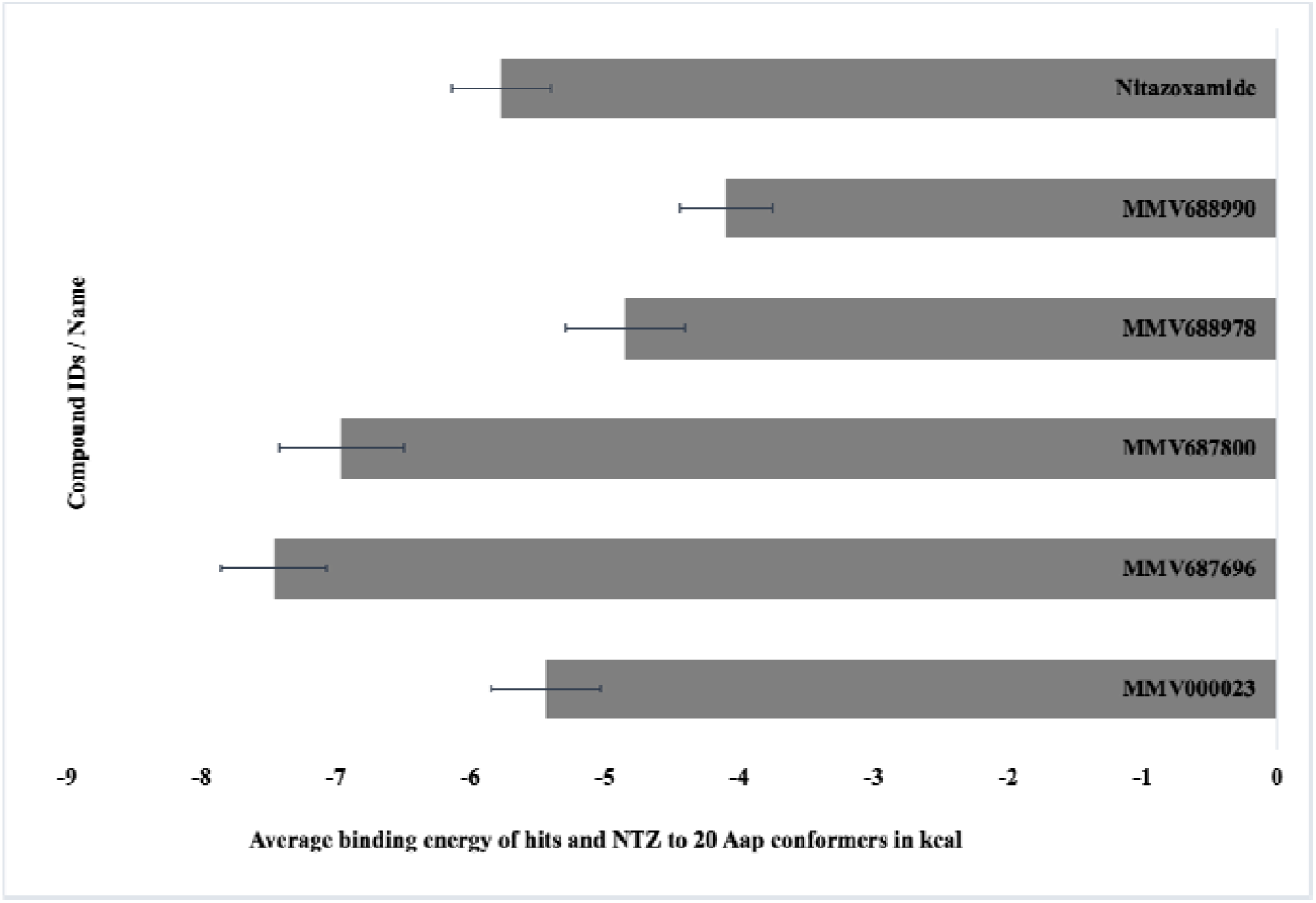
Average binding energy of 5 hits and NTZ to 20 conformers of Aap (kcal/mol). M7MV687696 and MMV687800 demonstrated the highest binding affinity for the 20 Aap conformers.

Since data from the isogenic 042 mutant biofilm inhibition assay additionally suggests that MMV687800 could have a second target (likely AAF/II), all five investigated hits were docked against AAF/ II major and minor subunits. The five hits and NTZ demonstrated different degrees of moderately thermodynamically favourable binding to the EAEC (AAF/II major and minor subunits), the only EAEC adhesins whose solution structure have been resolved [8]. Hits demonstrated stronger binding to non-Gd site cavities present on the surfaces of the EAEC proteins (Table 4). NTZ, initially shown to inhibit aggregative adherence fimbria and type I pili assembled by the Chaperone Usher (CU) pathway in EAEC was subsequently proven to interfere with the folding of the usher beta-barrel domain in the outer membrane [45]. It was recently reported for its selective activity in disrupting beta-barrel assembly machine (BAM)-mediated folding of the outer membrane usher protein in uropathogenic *Escherichia. coli* (UPEC) [46]). In our screening it demonstrated the highest affinity (−6.6 kcal/mol) for AAF/II closely followed by MMV687800 with ΔG of -5.7 kcal/mol. With focused binding interaction (that is, focusing on the Gd site), the strongest binding interaction with AAF/II major subunit among the hits was obtained with MMV687800 (−5.7 kcal/ mol), MMV000023 (−5.5 kcal/mol), then MMV688978 (−5.2 kcal/mol) (Table 4). In all, Nitazoxanide, reported to interfere with AAF/ II assembly [28], outperformed all five test compounds with respect to the strength of interaction with the AAF /II binding site even though the differences are at best marginal.

**Table 4:** Free energy of interaction between hit molecules and AAF/II major and minor subunits of EAEC strain 042.

| Hits / NTZ | AAF/II major subunit binding free energy (kcal/mol) |  | AAF/II minor subunit binding free energy (kcal/mol) |  |
| --- | --- | --- | --- | --- |
|  | Blind docking | Focused docking | Blind docking | Focused docking |
| MMV000023 | -5.6 | -5.5 | -6.7 | -4.7 |
| MMV687696 | -8.6 | -3.8 | -9.6 | -4.1 |
| MMV687800 | -6.9 | -5.7 | -8.8 | -4.4 |
| MMV688978 | -5.0 | -5.2 | -6.3 | -4.2 |
| MMV688990 | -4.7 | -4.5 | -5.0 | -3.9 |
| Nitazoxanide | -5.4 | -6.5 | -6.2 | -5.5 |

## Discussion

Biofilms contribute significantly to pathogenesis of several bacterial infections [47-51]. They are major players thwarting host defense and the activity of antimicrobials against many microorganisms [52] and will consequently require a novel approach to target them during infections. EAEC are known to form copious biofilms, which enhances their persistence during infection [53,54] hence antibiofilm agents could be particularly effective [55]. As antibiofilm agents attack a colonization process and not bacterial viability, it is hoped that, unlike conventional antibiotics, they will exert less selection pressure for antimicrobial resistance [56].

Whilst antibiofilm activity of a few synthetic and natural small molecules has been demonstrated *in vitro* [52,57-62] and *in vivo* [55,63,64], for other pathogens, only one study reported significant antibiofilm activity of a compound, nitazoxanide, against EAEC biofilms [28]. Shamir *et al* (2010) discovered the antibiofilm activity of NTZ when they were evaluating its growth inhibitory activity and subsequently showed that this antiparasitic compound inhibits assembly of AAF/II. NTZ has subsequently been shown to interfere with pilus usher function [45,46] and Bolick *et al* (2013) [55] were able to show that it reduces diarrhoea *in vivo*, and shedding of EAEC at concentrations not inhibiting growth. These data suggest that antiadhesive agents have therapeutic potential against EAEC. By systematically screening for EAEC antibiofilm activity, this study has uncovered many more inhibitors that are significantly more potent than NTZ. At least one of them, MMV687800 inhibits EAEC-specific targets and so is unlikely to have deleterious effects on the normal flora.

The five hits we identified from the MMV’s pathogen box reproducibly inhibited biofilm formation by one or two EAEC strains by 30-85% while inhibiting growth by ≤ 10 at 5 µM. Our hit rate for biofilm inhibitors that do not inhibit growth (1.25%), from a curated library, provides support for this drug discovery approach. Targeting EAEC biofilms without compromising their viability holds a high potential since hits from our screen are unlikely to select for antibacterial resistance due to selection pressure for resistant strains.

We report here antibiofilm activity of five compounds, most of which are not new to the drug industry thus providing a good starting point for antibiofilm drug discovery. Four of our hits MMV687800, MMV688978, MMV000023, and MMV688990 are existing therapies for other neglected tropical diseases, but not diarrhoea or Gram-negative infections. Clofazimine (MMV687800) is used to treat leprosy (*Mycobacterium leprae)*: https://www.drugbank.ca/drugs/DB00845 and its antimicrobial spectrum does not include Gram negative organisms like EAEC [65,66]. The anti-tubercular drug was recently reported to demonstrate a broad activity spectrum against coronaviruses including SARS-CoV-2 [67]. Miltefosine (MMV688990) is an anti-leishmanial drug [68] originally developed as an anti-cancer drug in 1980s: https://www.drugbank.ca/drugs/DB09031, Auranofin (MMV688978) is a gold salt used in treating rheumatoid arthritis: https://www.drugbank.ca/drugs/DB00995 but is now been tried for several other disease conditions, not including diarrhoea [69]. Primaquine (MMV000023) is an antimalarial agent: https://www.drugbank.ca/drugs/DB01087, which is active against gametocytes as well as recalcitrant parasitic forms of malaria, and therefore useful for preventing vivax and ovale relapse [70]. None of these four compounds has any previously documented antibacterial activity (*in vivo* or *in vitro*) against Gram negative bacteria, however, Ugboko *et al* (2019) in an *in silico* screen and analysis of broad-spectrum molecular targets and lead compounds for potential anti-diarrhoea agents reported MMV687800 as one of the compounds with predicted activity against potential targets for diarrhoea disease among twenty five other compounds [71]. Low *et al* (2017) in screen to identify TB active compounds against nontuberculous Mycobacterium additionally retrieved MMV687800 as an *M. avium*-specific main hits among five other MMV compounds [72] and Hennessey *et al* (2018) also reported MMV687800 among seven MMV compounds as an inhibitor with dual efficacy against *Giardia lamblia* and *Cryptosporidium parvum* in a pathogen box screen [73].

Rollin-Pinheiro *et al* (2021) in a screen to identify antifungal drugs against *Scedosporium* and *Lomentospora* species from the pathogen box chemical library retrieved MMV688978 as hit in addition to discovering its ability to decreasee the fungal biomass of preformed biofilm by about 50% of at 1 × MIC for *S. aurantiacum* and 70% at 4 × MIC of *S. dehoogii* and *L. prolificans* [74]. MMV687696, has been reported by MMV to possess anti-tuberculosis activity.

The antiparasitic compound nitazoxanide, also a component of Pathogen Box, which was previously reported to inhibit biofilm by EAEC 042 [28] was not recovered as hit from our screen and we verified that it is indeed inactive at 5 µM, our screen concentration. We re-tested nitazoxanide at concentrations for which antibiofilm activity was previously recorded and observed significant concentration-dependent biofilm inhibition at 15, 20 and 25 µg/ml (48.8, 65.15 and 81.4 µM) but with estimated growth inhibition of up to 50% [28]. Consequently, the validated hits obtained from the screen in this study, which lack growth inhibition activity at the concentration tested, were at least 10 times more active in biofilm inhibition than nitazoxanide the only known EAEC biofilm inhibitor, which has subsequently been shown to be effective against experimental infections in a weaned mouse model [55]. Unlike NTZ, our five hits inhibited biofilm formation at concentrations that did not produce significant growth inhibition, pointing to the possibility of biofilm-specific targets and minimal, if any, cross resistance with clinical antibacterials [44,56]. For three of the compounds for which we could get sufficient chemical to test, inhibition was largely concentration dependent, again suggesting that specific biofilm factors are targeted. When those factors are EAEC-specific, use of antibiofilm agents is unlikely to disrupt the normal flora, including other *E. coli*, which are protective against enteric infection. Biofilm formation is a complex and stepwise process involving numerous bacterial factors, which vary among and even within pathotypes. Early-stage contributors include adhesins, flagella and secreted protein autotransporters. At late-exponential phase, the accumulation of quorum sensing signals leads to the activation of other genes. Late-stage biofilm factors include adhesins with greater permanence, components that comprise or requite a macromolecular matrix as well as antiagregation proteins which can be co-opted to release bacteria from the biofilm.

The temporal patterns of biofilm inhibition and EAEC inhibition spectra of compounds MMV687800, MMV688978, MMV687696, MMV000023 and MMV688990 were different, implying that they likely target different contributors to EAEC biofilm formation. EAEC expressed many surface factors involved in host adherence and biofilm formation [21,40,75.76]. Any of these, or their regulators, could be direct targets. Overlaying activity spectra data with virulence factor profiles of genome sequenced EAEC strains provided preliminary insights to mechanism of action. For MMV687800, subsequent testing of five isogenic mutants of EAEC 042 provided further insight. Aap is an antiaggregation protein or dispersin that allows bacteria to detach from old biofilms and seed new ones. Mutants in *aap* show increased biofilm formation but impaired colonization [21,40]. Biofilm formation by *aap* mutant (LV1: 042Δ*aap*) was significantly less inhibited by MMV687800 than wildtype. Additionally (LV2: 042Δ*aap*Δ*hra1*) double mutants [40] were inhibited in biofilm formation to a proportionally lower degree and inhibition seen in (LTW1: 042Δ*aap*Δ*aaf*) was statistically insignificant compared to controls no compound at all (p = 0.11). This phenotype could be complemented in trans and thus molecular Koch’s postulates [77] are fulfilled for *aap* as an MMV687800 target.

To preliminarily determine whether the interaction between MMV687800 and Aap could be direct, we independently determined binding affinities of hits with twenty Aap conformers using molecular docking techniques since the solution structure of Aap has been resolved by NMR [35]. The strong binding interaction sustained for hit MMV687800 and in the outcome of the molecular computational docking experiments against 20 conformational instances of the dispersin (Aap) in Figure 7 indicates a high significance for the obtained affinities. MMV687800 demonstrated moderately strong interaction (in some cases ΔG ≤ -8.1 kcal) with Aap. This additionally suggests that *aap* is one of the surface factors targeted by MMV687800.

The *aafA* gene encodes the structural subunit of AAF/II fimbriae and Hra1, the heat-resistant agglutinin 1 is an outer-membrane protein involved in autoaggregation that serves as an accessory colonization factor [40,75,78]. MMV687800 showed a small, insignificant reduction of activity in the *aafA* mutant 3.4.14 [79], but not the *hra1* mutant, which we initially discounted, given the robust phenotype with Aap. However, the reduction in MMV687800 biofilm inhibition was visible with the *aap aafA* mutant LTW1, and complementation of this mutant with *aap* alone LTW1(pDAK24) could not fully restore biofilm inhibition. These data, and the comparative genomic data which shows that *aafA-*D are unique to biofilm inhibition groups alone (data not shown), strengthen the likelihood that AAF/II is implicated in MMV687800 biofilm inhibition, albeit to a lower degree than Aap. The five hit compounds demonstrated stronger binding to non-Gd site cavities present on the surfaces of AAF/II (Table 4). Docking outcome however revealed NTZ, known to inhibit AAF/II assembly exhibited highest affinity (−6.6 kcal/mol) for AAF/II followed by MMV687800 which had -5.7 kcal/mol again, indicating that AAF/II could be a likely target for MMV687800.

This study has some limitations. EAEC are highly heterogeneous and the full spectrum of lineages and virulence factors is not covered by the two strains used for this screen, or even by the 27 strains employed to better understand activity profiles of the hits. We determined the probable mechanism of action of only one hit, taking advantage of the easily-generated VirulenceFinder output and the bank of mutants we had on hand. In doing so, we were able to generate proof of principle and rule out Aap, AAF/II and Hra1 as targets for the other compounds. However, it is conceivable that at least some of the targets of the other compounds may not be unique to EAEC or may be genes of unknown function. More intensive and unbiased comparative genomic approaches, currently underway, will be needed to exhaustively screen for the targets of the other four hits, which could well be more promising than the hit highlighted in this study.

In conclusion, this study identified five biofilm inhibiting but non-growth inhibiting compounds that have not been previously described as bacterial anti-adhesins or Gram-negative antibacterials. Hits discovered from this screen will add to the drug discovery pipeline for this neglected pathogen, improve understanding on EAEC colonization and enhance EAEC-based interventions. Additionally, the understandings from our experiments that some of our hits are unlikely to target known EAEC adhesins is initiating investigations at the molecular level which can open us up to a world of novel adhesins and consequently enhance our understanding of EAEC pathogenic factors.

## Acknowledgement

The authors are grateful to the following people for technical assistance during the project: Jeremiah J. Oloche, Olabisi C. Akinlabi, Jesuferanmi Igbinigie, Erkison Ewazimo Odih, Anderson O Oaikhena, Taiwo Badejo, Adewole D Pelumi, Stella Ekpo, Abiodun Oyerinde, Catherine Oladipo, Emmanuel Bamidele, El-Shama Q Nwoko, Amos Olowokere, Uchechi Okoroafor and Joy Olorundare.

We thank the Medicines for Malaria Venture (MMV) for supplying Pathogen Box, supplementary amounts of select chemicals from the box, as well as for drug discovery training for DAK. We additionally thank the Wellcome Trust for advanced training in drug discovery for DAK.

## Author Contributions

**Conceptualization:** Iruka N. Okeke.

**Data curation:** David A. Kwasi, Chinedum P. Babalola, Olujide O. Olubiyi and Iruka N. Okeke.

**Formal analysis:** David A. Kwasi, Olujide O. Olubiyi and Iruka N. Okeke.

**Investigation:** David A. Kwasi, Jennifer Hoffmann, Olujide O. Olubiyi and Iruka N. Okeke.

**Methodology:** David A. Kwasi, Jennifer Hoffmann, Olujide O. Olubiyi, Ikemefuna C. Uzochukwu and Iruka N. Okeke.

**Resources:** Iruka N Okeke, Olujide O. Olubiyi

**Supervision:** Olujide O. Olubiyi, Chinedum P. Babalola and Iruka N. Okeke

**Validation:** David A. Kwasi, Chinedum P. Babalola, Ikemefuna C. Uzochukwu, Olujide O. Olubiyi and Iruka N. Okeke.

**Writing – Drafts:** David A. Kwasi, Chinedum P. Babalola, Olujide O. Olubiyi, Ikemefuna C. Uzochukwu and Iruka N. Okeke.

## Supplementary data

**Table S1:**
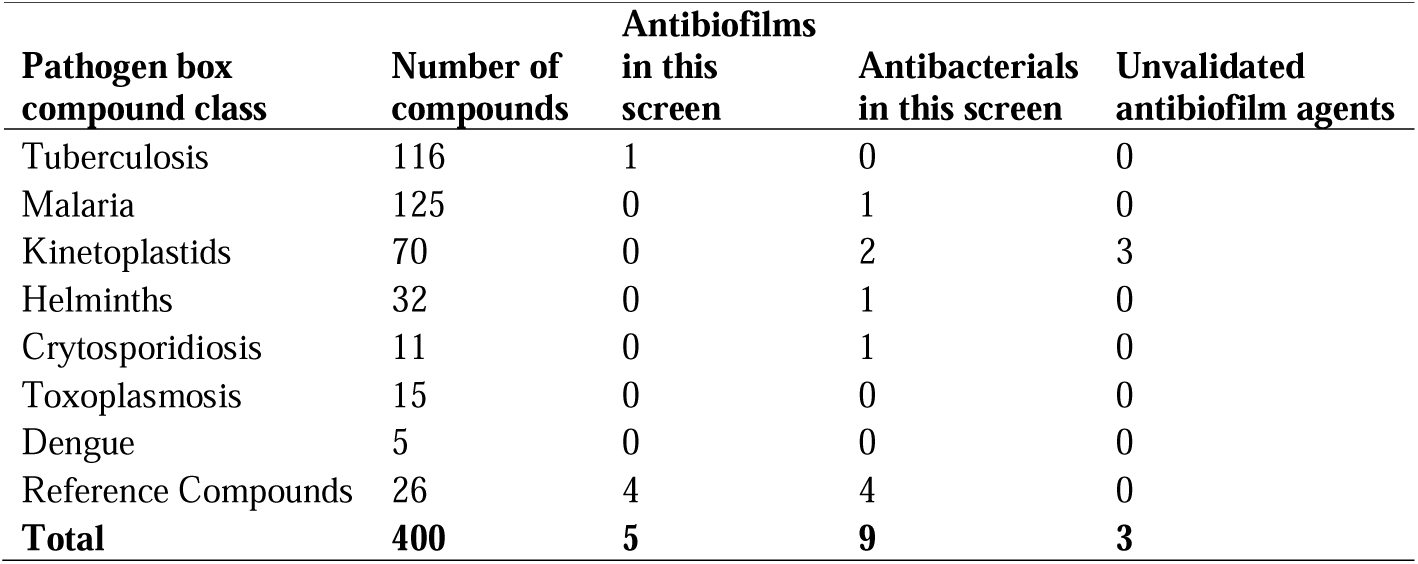
Classification of pathogen box compounds matched with biofilm inhibition screen outcome.

**Figure S1:**
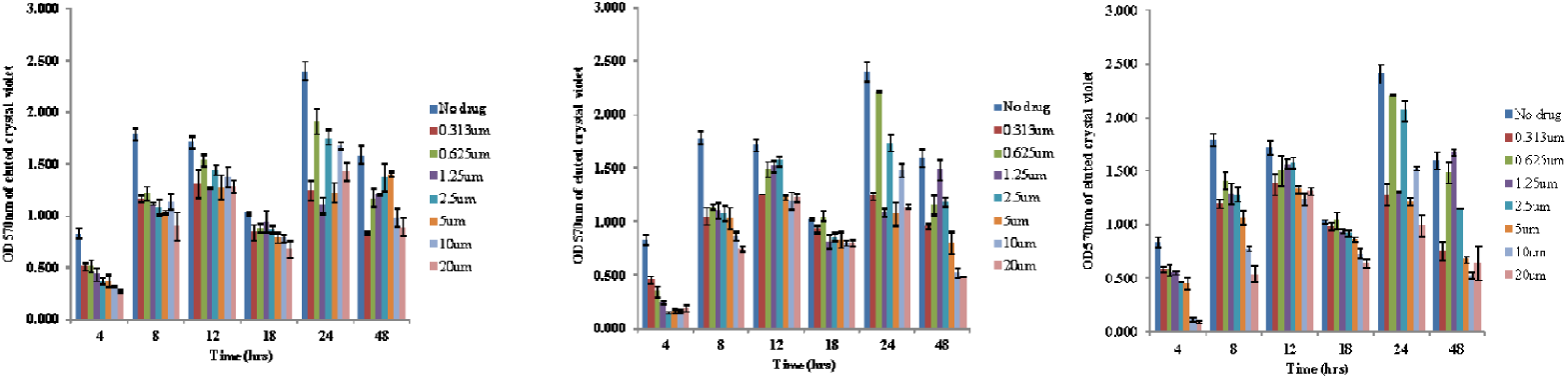
Time course inhibition of biofilms in EAEC 042 using different hit concentrations of (a) MMV688978, (b) MMV687800, (c) MMV688990.

